## Supporting Information for "Object representations in the human brain reflect the co-occurrence statistics of vision and language"

SI figures for:

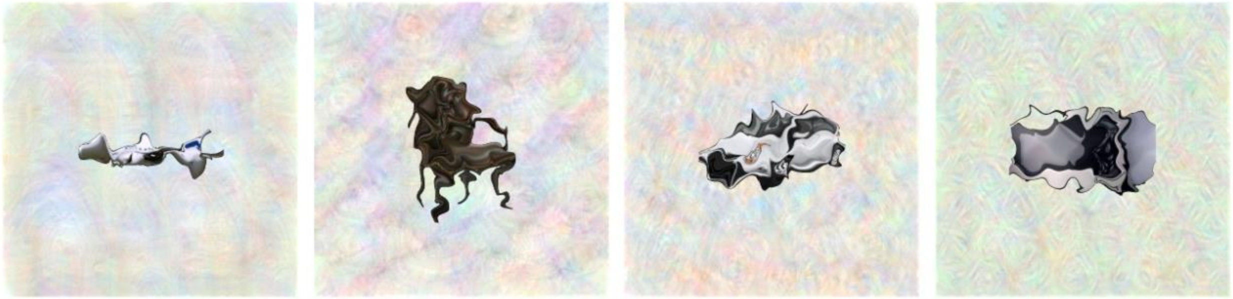

***Figure S1. Warped object stimuli.*** *In the fMRI experiment, subjects were asked to press a button whenever they saw an image of a warped object. These warped objects were created by applying a diffeomorphic warping algorithm to the images in our stimulus set. This figure shows examples of warped images for four different categories of objects.*

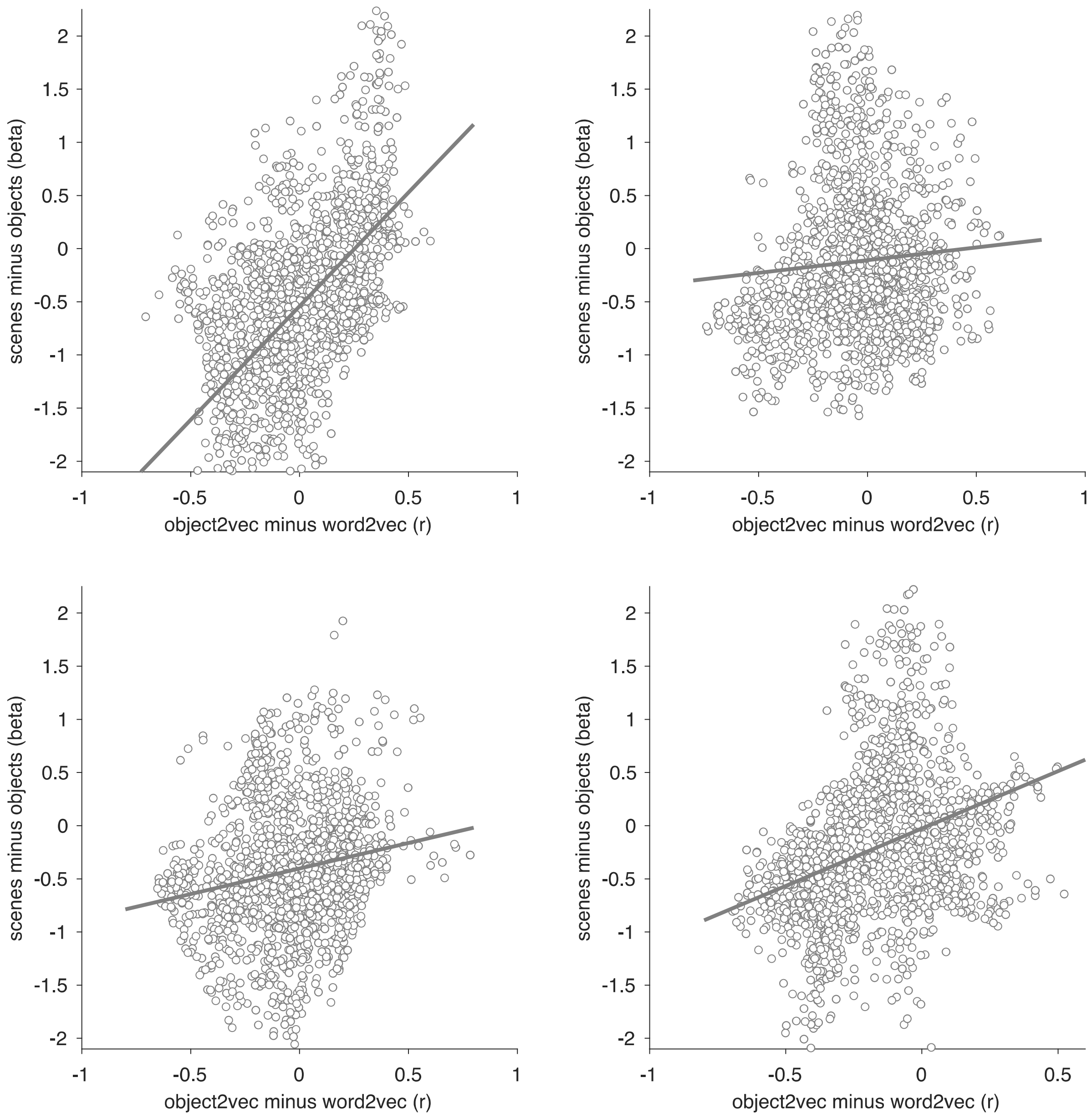


***Figure S2. Relationship between encoding-model accuracy and category selectivity in single subjects.*** *These scatter plots show how the voxelwise encoding-model effects are related to category selectivity within each subject. Each point is a voxel, and these plots include all voxels with significant effects for either the image-based object2vec encoding model or the language-based word2vec encoding model. The x-axis plots the difference in prediction accuracy between the object2vec encoding model and the word2vec encoding model. The y-axis plots the difference in activation between scenes and objects, based on data from a separate set of functional localizer runs. Positive linear trends suggest that voxels that are better predicted by object2vec than word2vec tend to be more scene-selective, whereas voxels that are better predicted by word2vec than object2vec tend to be more object-selective (clockwise from top left, Pearson r values = 0.55, 0.07, 0.19, 0.32).*

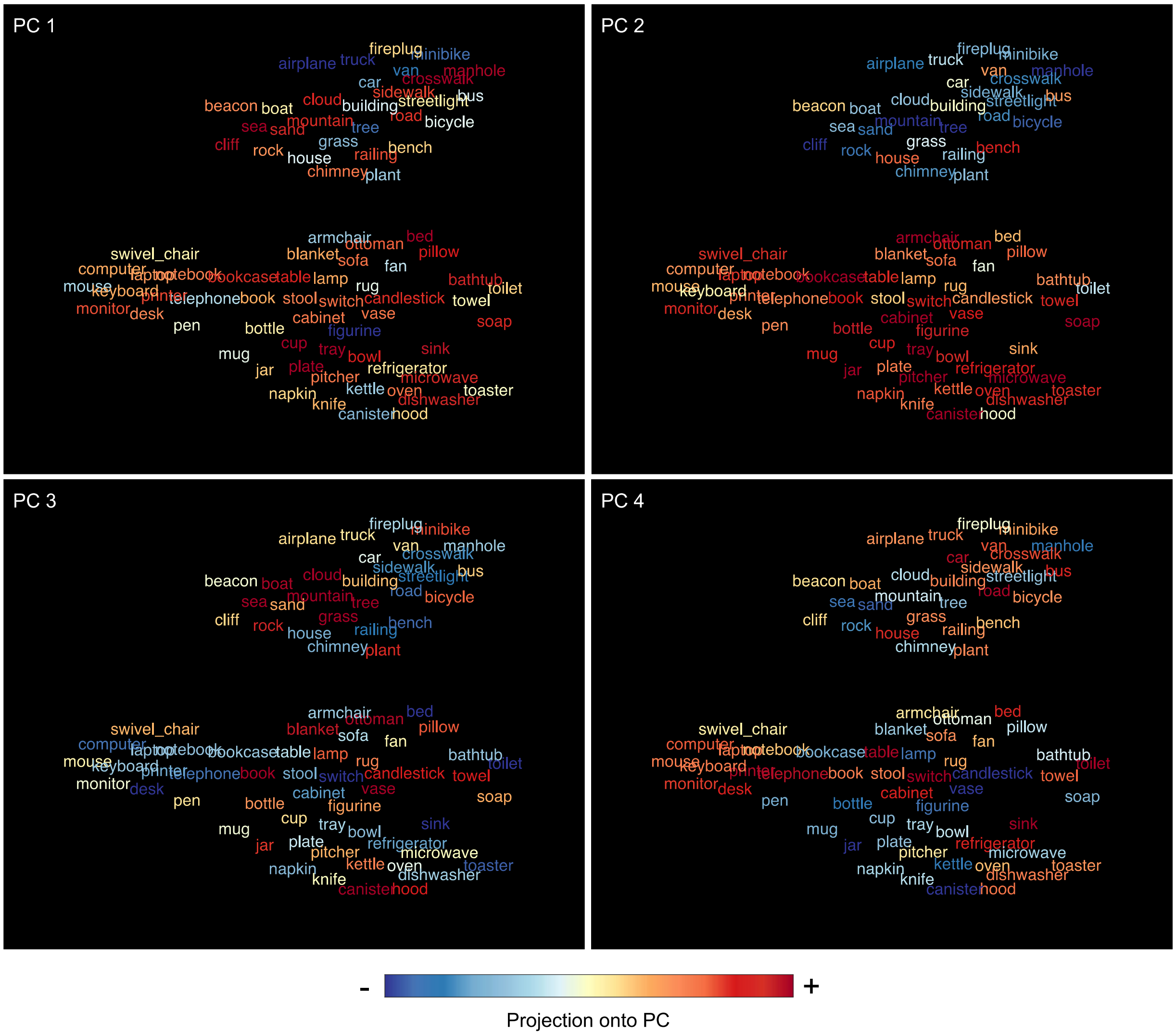


***Figure S3. Principal components of voxel tuning for language-based object context.*** *Principal components analysis was used to examine variance in encoding-model regression weights across voxels. This plot illustrates the first four principal components (PCs) of the regression weights for the language-based word2vec encoding model, using all voxels with significant prediction accuracies. The 81 object categories from the fMRI experiment were projected onto each PC (as indicated by the color coding from blue to red). The spatial arrangement of the object categories is the same as the tSNE plot in Figure 1.*

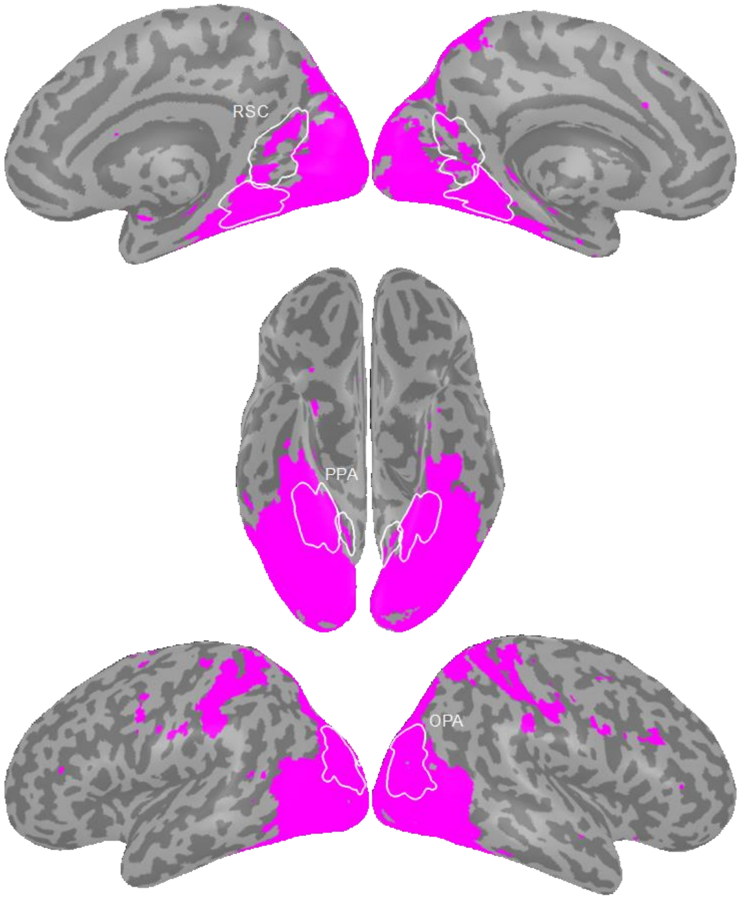


***Figure S4. Split-half reliability mask.*** *Encoding model analyses were performed for all voxels with split-half reliability scores greater than or equal to r = 0.1841, which corresponds to a p-value of 0.05. For each voxel, split-half reliability was calculated as the Pearson correlation between the mean responses to the 81 object categories in odd and even runs. The purple regions on this cortical surface rendering indicate voxels that surpassed the reliability threshold. ROI parcels are shown for scene-selective ROIs. PPA = parahippocampal place area, OPA = occipital place area, RSC = retrosplenial complex.*
